# Low-intensity transcranial focused ultrasound engages parvalbumin-positive GABAergic interneurons in a humanized mouse model of chronic pain: from electrophysiology to cellular investigation

**DOI:** 10.1101/2025.10.08.680737

**Authors:** Min Gon Kim, Chih-Yu Yeh, Huan Gao, Keunhyung Lee, Kalpna Gupta, Bin He

**Affiliations:** Department of Biomedical Engineering, Carnegie Mellon University, Pittsburgh, PA 15213, USA; Department of Medicine, University of California at Irvine, Irvine, CA 92697, USA; Division of Hematology, Oncology and Transplantation, University of Minnesota, Minneapolis, MN 55455, USA; Neuroscience Institute, Carnegie Mellon University, Pittsburgh, PA 15213, USA

**Author notes:** These authors contributed equally to this work (Min Gon Kim, Chih-Yu Yeh). Corresponding Author: Bin He, Ph.D., Department of Biomedical Engineering Carnegie Mellon University 5000 Forbes Avenue, Pittsburgh, PA 15213, USA.

## Abstract

**Background:** Low-intensity transcranial focused ultrasound (tFUS) offers high spatial specificity and deep brain penetration, showing great promise as a non-invasive stimulation technology for modulating brain activity and behavior. Recent studies show that specific tFUS parameters targeted to pain-processing brain circuits can significantly alter pain-related behaviors in rodent models and humans. However, a comprehensive understanding of how tFUS influences brain networks and cellular mechanisms is essential to optimize efficacy and facilitate safe translation to clinical pain therapies.

**Objective:** We aimed to evaluate the modulation of inhibitory neural circuits induced by tFUS of 40 Hz pulse repetition frequency (PRF) in a humanized mouse model of chronic pain, integrating local and network-level electrophysiological investigations, molecular analyses, and histological assessment to confirm safety.

**Methods and Results:** We used a 128-element random array transducer for stimulation, along with a non-invasive and flexible 30-channel electroencephalography (EEG) to assess local evoked responses, topographical brain activity, and global brain dynamics including excitation and inhibition (E/I) balance. To further assess tFUS neuromodulation effects at the cellular level, we performed immunohistochemistry (IHC) analysis and found that tFUS significantly increased the activity of inhibitory neurons as indicated by elevated expression of Glutamate Decarboxylase 67 (GAD67) and Parvalbumin (PV). Finally, safety was evaluated in the same brain samples used for mechanistic analysis, with blinded histological assessment revealing no signs of tissue damage.

**Conclusions:** These findings provide new evidence that tFUS non-invasively engages PV GABAergic inhibitory circuits in a chronic pain mouse model, supporting its development as a robust neuromodulation strategy.

**Topics:** Chronic pain; Transcranial focused ultrasound; Non-invasive brain neuromodulation; GABAergic neural circuit modulation

**Highlights:**

- Multi-modal assessment of low-intensity tFUS in a humanized chronic pain model.
- 40 Hz tFUS enhances inhibition, mirroring optogenetic PV⁺ neuron activation.
- Repeated tFUS restored chronic pain-disrupted E/I balance.
- Multi-session tFUS upregulates GAD67 and PV⁺ interneuron expressions.

## 1. Introduction

Treating chronic pain by means of non-pharmacological approaches is urgently needed due to the severe side effects of drugs including abuse, overdose, and addiction, all of which contribute to a significant health pandemic [1–3]. To reduce reliance on pharmacological treatments, neuromodulation strategies targeting pain-processing circuits have been developed [4–9]. Deep brain stimulation (DBS) is effective for movement disorders and has shown promise in managing chronic pain [4,5], but it requires invasive surgical implantation [4,6,8]. Non-invasive neuromodulatory alternatives such as transcranial direct current stimulation (tDCS) and transcranial magnetic stimulation (TMS) demonstrated diverse treatment efficacies [10–13]; however, they typically influence broad areas and have limited ability to reach deeper brain structures [6,7].

Transcranial focused ultrasound (tFUS) is a promising neuromodulation technology with high spatial specificity and penetration depth compared to other non-surgical brain stimulation methods [14–17], enabling non-invasive modulation of targeted brain regions [14–19] along with behavior control [20,21] and treatment of neuropsychological disorders [22–26]. Low-intensity tFUS has recently shown potential for pain treatment in both pre-clinical and human studies. In humans, tFUS targeting the anterior cingulate cortex modulated local activity [27] and alleviated both chronic pain symptoms and responses to acute heat stimuli [28]. Additional evidence demonstrated antinociceptive effects from anterior thalamus stimulation [29] and mood improvements with posterior frontal cortex tFUS in pain patients [30]. In animal models, stimulation of the periaqueductal gray (PAG) has been shown to attenuate formalin-induced nociceptive activity in the dorsal horn of the spinal cord [31], while stimulation of the primary somatosensory cortex (S1), insula, or both regions simultaneously significantly suppressed pain-related behaviors in a humanized sickle cell disease (SCD) mouse model [32,33]. These studies suggest that targeted neuromodulation in the brain can ameliorate pain; however, further investigation is needed to elucidate the underlying mechanisms driving the modulation of pain-related behaviors and neural circuits. A deeper understanding of both cellular and electrophysiological effects, including sustained modulation of neuronal expression and network-level dynamics, will be essential to bridge critical knowledge gaps.

In this study, we investigated the mechanisms and safety of low-intensity tFUS via integrated electrophysiological and cellular approaches in a humanized mouse model of chronic pain. We hypothesize that tFUS with a lower pulse repetition frequency (PRF) can significantly activate specific GABAergic interneurons, such as the Parvalbumin (PV) cells, and enhance inhibitory effects in the mouse model of SCD which expresses >99% human sickle hemoglobin and exhibits both severe chronic pain and hyperalgesia [34,35]. We first examined local evoked responses, topographical brain activity, global network dynamics in excitation/inhibition (E/I) balance using flexible 30-channel electroencephalography (EEG) recordings paired with a highly-focused 128-element random array transducer. At the cellular level, we next incorporated immunohistochemistry (IHC) targeting inhibitory circuit markers, including PV and Glutamate Decarboxylase 67 (GAD67), a key GABA-synthesizing enzyme [36–39]. Additionally, we evaluated the safety profile of 14-day repeated tFUS through blinded histological analysis of the same brain samples examined by IHC. This comprehensive approach aimed to uncover how low-intensity tFUS non-invasively modulates the inhibitory circuits at both the cellular and systems levels, thereby providing foundational insights into its translational potential for chronic pain.

## 2. Materials and methods

### 2.1 Animals

All animal experimental procedures were approved by the Institutional Animal Care and Use Committee of Carnegie Mellon University and the Long Beach VA Medical Center, and complied with the National Institutes of Health guidelines. Female wild-type mice (C57BL/6J, Jackson Laboratory, Bar Harbor, ME, USA), female and male humanized transgenic homozygous Berkley sickle cell mice (HbSS-BERK), and male Wistar outbred rats (Hsd:WI, Envigo Laboratory, Indianapolis, IN, USA) were used as subjects. HbSS-BERK mice were generated in-house by cross-breeding homozygous (SS) males with hemizygous (AS) females [40]. (For detailed mice information, see Supplementary Method S1.) Older mice were used in this study, including HbSS-BERK mice (13-17 months) and age-matched wild-type mice (11-14 months), as the hallmark feature chronic pain severity in SCD is known to worsen with age [33,35,41].

### 2.2 Characterization of custom-developed 128-element random array transducer

To achieve high spatial specificity for precise stimulation within the small mouse brain, a previously developed spherically-concave and highly-focused 128-element random array ultrasound transducer at 1.5 MHz center frequency (H276, Sonic Concepts, Inc., Bothell, WA, USA) was used in this study [32].

To evaluate the pressure and intensity profiles of tFUS through the mouse skull, we utilized a customized *ex-vivo* 3D acoustic field mapping system with a needle hydrophone (HNR500, Onda Corporation, Sunnyvale, CA, USA) and a three-axis positioning stage (XSlide, Velmex, Inc., Bloomfield, NY, USA) [33,42]. A freshly harvested skull from a euthanized mouse was positioned between the array transducer and the hydrophone to closely mimic the *in vivo* tFUS configuration.

### 2.3 tFUS stimulation targeting procedures and parameters

Mice were anesthetized using 3% isoflurane with a 1 L/min oxygen flow for induction, and 1-1.5% isoflurane with a 0.3-0.5 L/min oxygen flow for maintenance. A heating pad was used to maintain body temperature, and ophthalmic ointment was applied to protect the eyes. Core body temperature was regularly monitored using a rectal temperature probe.

To estimate the targeted brain circuits using a 128-element random array transducer, we followed previously established targeting procedures [32]. Briefly, the Allen Mouse Brain Atlas was referenced to determine the coordinate of the ultrasound focus. The transducer was positioned by visually aligning the transducer’s X-axis with the posterior edge of zygomatic structure and the Y-axis with the mouse’s rostral caudal midline (Fig. 1A-B). We applied single- or multi-session tFUS with specific parameters including tone-burst duration (TBD), ultrasound duration (UD), inter-sonication interval (ISoI), pulse repetition frequency (PRF), and total sonication time (Fig. 1C).

**Figure 1.**
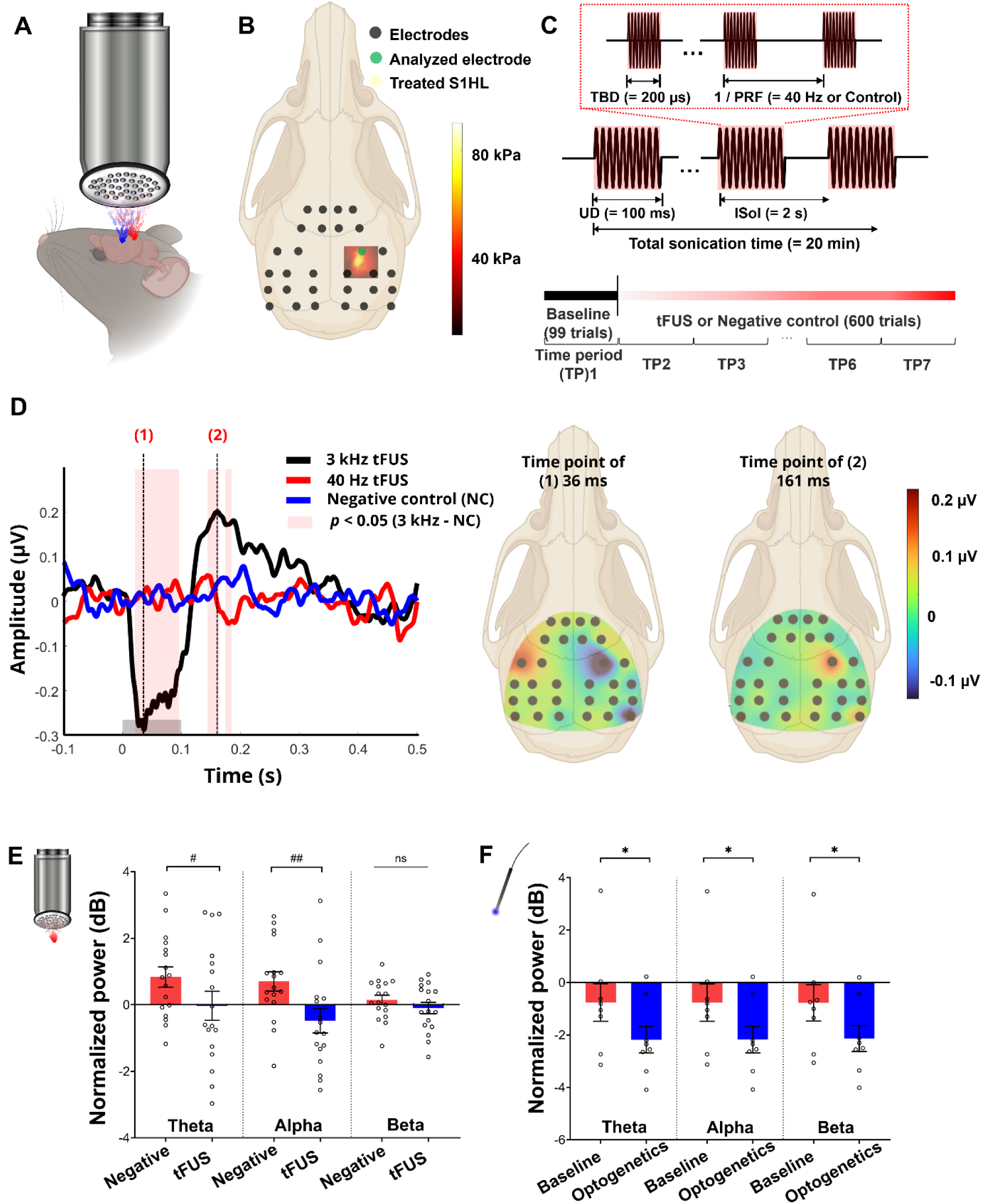
Single-session tFUS induced local brain circuit modulation revealed by EEG. (A) Schematic of tFUS stimulation using a 128-element random array transducer. It features dynamic beam steering capabilities as demonstrated by red and blue beams when targeting specific brain circuits. (B) EEG recording setup with 30-channel flexible electrode along with highly focused ultrasound stimulation beam as presented by S1HL stimulation with the location of the analyzed electrode representing S1HL activity. (C) Single-session tFUS stimulation with selected parameters: a PRF of 40 Hz or control value, a UD of 100-ms, an ISoI of 2-sec, and a total sonication time of 20-min. Recording timeline below illustrates EEG acquisition during a 99-trial baseline period (TP1) followed by 600 trials of tFUS stimulation or control (TP2-TP7). (D) Representative EEG traces and topographic maps during tFUS with PRF of a 3 kHz related to excitatory effect show that a single-session tFUS can perturb local brain activity. The gray rectangle represents the tFUS stimulation or negative control period. (E) Significant suppression of low-frequency EEG oscillations, particularly within the theta and alpha bands, was observed with 40 Hz PRF tFUS in SCD mice (n=17). (F) Significant suppression of low-frequency EEG oscillations, especially in the theta, alpha, and beta bands, was similarly observed with optogenetic activation of PV-positive interneurons in rats (n=8). ^#^*p* <0.05, ^##^*p* <0.01 using *t* test with Mann-Whitney test; \**p* <0.05 using *t* test with Wilcoxon matched-pairs signed rank test. ns, not significant.

### 2.4 *In vivo* electrophysiology recordings and data analysis in mice

After securing the mouse head, cranial fur was removed using a depilatory cream under the isoflurane anesthesia. EEG recordings were acquired using a 30-channel mouse electrode array (HC32 Mouse EEG, NeuroNexus, Ann Arbor, MI, USA) placed on the scalp and stabilized with saline. To analyze neuronal oscillations during resting-state activity and intrinsic response to tFUS, EEG signals were obtained from the electrode positioned directly above and closest to the hindlimb region of the primary somatosensory cortex (S1HL), identified using anatomical landmarks (Fig. 1B). Although scalp EEG is influenced by volume conduction, it reliably represents cortical electrical dynamics [43]. EEG data acquisition was performed through a ZIF-Clip headstage connected to a PZ5 NeuroDigitizer and Synapse software (Tucker-Davis Technologies, Alachua, FL, USA). Data was preprocessed using TDT-provided offline scripts and subsequently analyzed with custom MATLAB scripts. Analyses included power spectral estimation, topographical mapping, and assessment of E/I balance indicated by aperiodic exponent [44,45]. (For detailed data analysis, see Supplementary Method S2.)

### 2.5 Histology, imaging, and analysis

After 14-days of tFUS stimulation or sham treatment, mice were euthanized, perfused, and brains were immediately collected and fixed following established protocol [32,46]. Paraffin-embedded sections (4 μm) were processed and immunostained at HistoWiz, Inc. (Long Island City, NY, USA) using an automated stainer (Leica Bond RX, Leica Microsystems, Nussloch, Germany) with primary antibodies included anti-GAD67, anti-GABA, PV, anti-CD31, and anti-Iba1, with detection performed using DAB reagents and hematoxylin. Hematoxylin and eosin (H&E) and terminal deoxynucleotidyl transferase dUTP nick end labeling (TUNEL) assays were also performed. Pathology assessment was blindly evaluated and confirmed by a senior board-certified research pathologist. (For detailed staining protocols and pathology assessment, see Supplementary Method S3.)

### 2.6 Virus injection and optical stimulation during recordings in rats

Wistar rats were anesthetized with isoflurane and secured in a stereotaxic apparatus. PV-ChR2(H134R)/cherry virus (Duke Viral Vector Core, Durham, NC, USA) was injected into left S1 identified with the rat brain atlas (1 μL at 1.5 mm depth) [47], using a Hamilton syringe and microinjection system. Animals were monitored following surgery, and recordings were performed 4–5 weeks later to allow viral expression. For neural recordings, a 32-channel EEG electrode (HC32 Rat Functional, NeuroNexus, Ann Arbor, MI, USA) was placed on the skull, and a 32-channel silicon optrode (A1×32-Poly3-10mm-50-177, NeuroNexus, Ann Arbor, MI, USA) was inserted into left S1. Pulses of blue light (465 nm, 50-ms, 2-sec interval) were delivered via PlexBright LED module (Plexon, Dallas, TX, USA) to selectively activate PV-positive interneurons expressing ChR2. (For detailed surgical and injection procedures, see Supplementary Method S4.)

### 2.7 Statistical analysis

Data were presented as mean ± standard error of the mean (s.e.m.). Statistical analysis was performed using commercial software (GraphPad Prism, San Diego, CA, USA). To assess changes relative to the pre-stimulation baseline and differences in brain responses within the same subject, a non-parametric paired t-test (Wilcoxon matched-pairs signed rank test) was employed with significance thresholds: ns: not significant; \**p*<0.05, \*\**p*<0.01, \*\*\**p*<0.001, \*\*\*\**p*<0.0001. Additionally, intergroup comparisons between different treatment conditions were evaluated using the non-parametric t-test (Mann-Whitney test) with significance levels: ns: not significant; ^#^*p*<0.05, ^##^*p*<0.01, ^###^*p*<0.001, ^####^*p*<0.0001.

## 3. RESULTS

### 3.1 Single-session tFUS suppressed local theta oscillation consistent with PV-driven inhibition

To investigate how specific tFUS parameters modulate intrinsic brain activity in pain circuits, we integrated 30-channel scalp EEG recordings with a 128-element random array ultrasound transducer to evaluate tFUS-evoked brain responses in SCD mice. We first tested the hypothesis that a single session tFUS could perturb local brain activity when precisely targeted to specific pain-processing brain regions. During EEG recordings, following a baseline period, ultrasound pulses were delivered to the right hemisphere’s S1HL region (200-µs TBD, 40 Hz or 3 kHz PRF, 100-ms UD, 2-sec ISoI, 20-min total sonication time; Fig. 1C). Using the excitation parameter with 3 kHz PRF, we found significant activation in the targeted S1HL compared with both the negative control and the 40 Hz PRF condition related to the inhibition parameter (Fig. 1D).

We next investigated whether intrinsic brain rhythms associated with chronic pain could be modulated by single-session tFUS with a PRF of 40 Hz, previously shown to suppress pain-related behaviors in SCD mice. We observed a significant suppression of low frequency brain oscillations, particularly in the theta and alpha bands during 40 Hz tFUS targeting S1HL region compared to the negative control (Fig. 1E). In contrast, 3 kHz tFUS produced excitatory effects, showing a bidirectional pattern when comparing the low versus high PRF responses relative to the negative control (Supplementary Fig. 1). To further investigate the mechanisms underlying tFUS neuromodulation, we performed optogenetic activation of PV-positive interneurons in rats, which significantly reduced theta, alpha, and beta band power (Fig. 1F), suggesting that PV activation suppresses low-frequency oscillations. These findings align with the previous report showing that silencing PV interneurons increases low-frequency activity [48], indicating that active PV interneurons constrain these rhythms and play a central inhibitory role in shaping cortical dynamics. The spectral outcomes of tFUS and optogenetics generally align yet not fully congruent, suggesting that tFUS may engage broader or complementary inhibitory networks compared to the specificity of optogenetic stimulation. Nevertheless, the shared suppression of the theta band suggests that the fast-spiking PV interneurons are key contributors to tFUS effects. While GABAergic interneurons such as PV and somatostatin-positive (SST) cells both influence theta oscillations, prior studies demonstrated that optogenetic suppression of SST reduces spontaneous low-frequency (<30 Hz) activity, whereas suppression of PV increases it [48,49], underscoring their distinct but complementary roles in modulating cortical oscillations. Thus, the observed suppression of theta power following tFUS likely reflect PV-driven inhibition rather than SST mechanisms.

### 3.2 Multi-session tFUS enhanced network-level inhibitory activity consistent with PV-mediated inhibition

In the local brain circuit, single-session tFUS stimulation with a PRF of 40 Hz resulted in a marked suppression of theta and alpha band power. However, oscillatory changes alone may not fully capture the complex dynamics of brain networks, particularly the balance between excitation and inhibition. We therefore analyzed the aperiodic exponent of EEG, computed by fitting the slope of the signal’s power spectrum as a marker of E/I balance (Fig. 2A). The E/I balance is a crucial aspect of neural circuit function, influencing the network’s ability to process information [45]. Shifts in this balance can indicate whether cortical activity becomes more excitatory reflected by a lower aperiodic exponent, or more inhibitory indicated by a higher aperiodic exponent [44,50,51]. We found a significant reduction in the aperiodic exponent in female SCD mice relative to female wild-type (Fig. 2B), reflecting a disrupted E/I balance commonly associated with chronic pain features.

**Figure 2.**
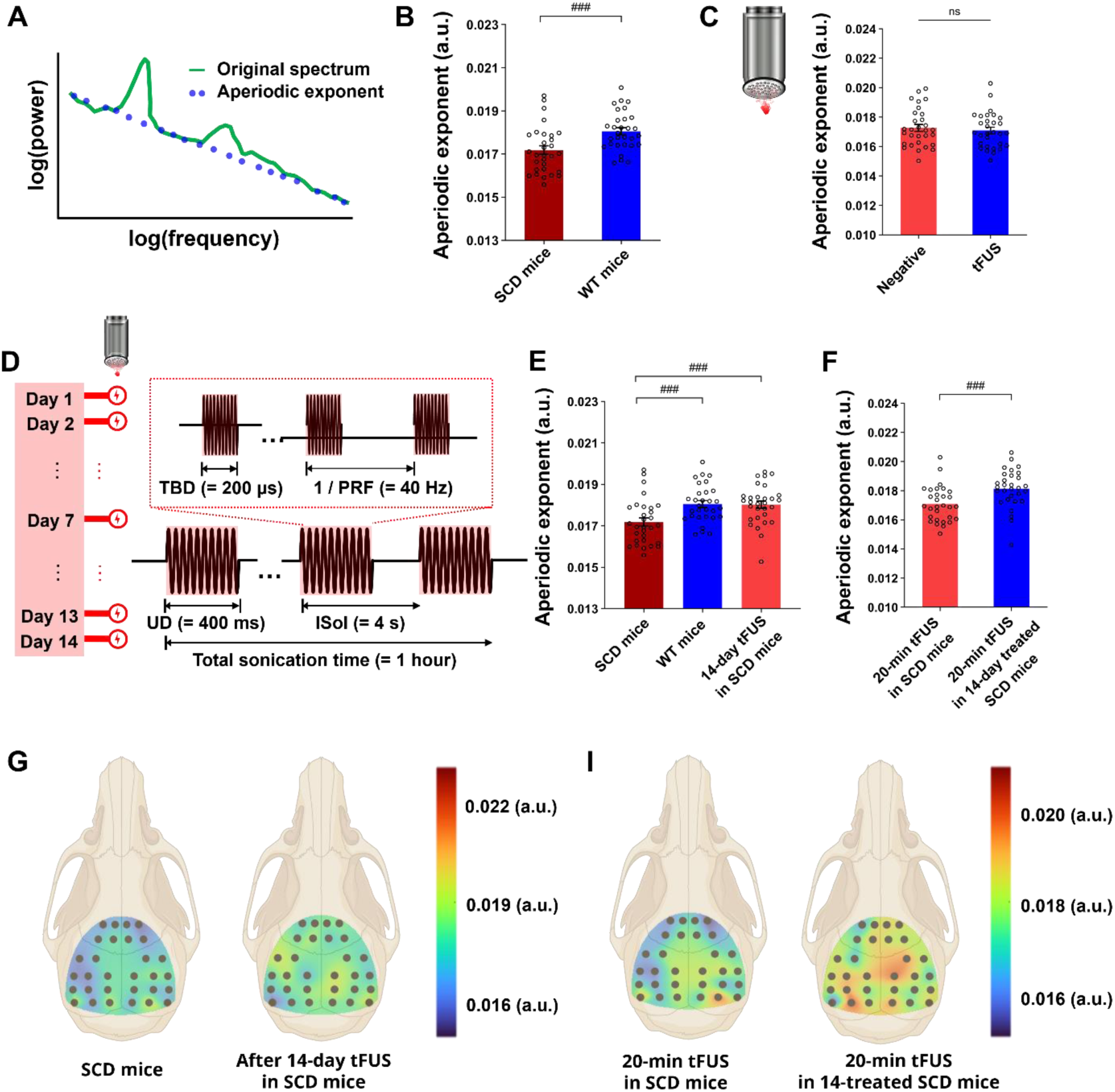
Multi-session tFUS induced global cortical inhibitory activity measured by EEG aperiodic exponent analysis. (A) Illustration of slope of the power spectrum showing the aperiodic exponent (blue) fit to the original power spectrum (green), used as a marker of E/I balance. (B) Aperiodic exponent was significantly reduced in SCD mice (n=17) compared to wild-type controls (n=8), suggesting a disrupted E/I balance. (C) Compared to negative control group, single-session tFUS with a PRF of 40 Hz did not produce a significant change in aperiodic exponent in SCD mice exhibiting chronic pain phenotypes. (D) Multi-session tFUS stimulation paradigm: 14 consecutive days of 1-hour daily tFUS stimulation with a PRF of 40 Hz, a UD of 400-ms, and an IsoI of 4-sec. (E) Aperiodic exponent significantly increased in SCD mice following multi-session tFUS at S1HL (n=5), reaching levels comparable to wild-type mice. (F) An additional single-session tFUS delivered after the 14-day stimulation significantly increased the aperiodic exponent compared to 20-min single-session tFUS in sickle mice, indicating amplified inhibitory effect. (G) Scalp EEG topographic maps of aperiodic exponent values showed a progressive increase from untreated SCD mice and post-multi-session tFUS. (I) Potential long-term synaptic plastic changes along with multi-session tFUS were investigated by comparing responses to single-session 20-min tFUS in SCD mice and an additional single-session tFUS in 14-day treated SCD mice. ^###^*p*<0.001 using *t* test with Mann-Whitney test; ns: not significant.

We then applied a single-session tFUS in female SCD mice; however, it did not significantly alter E/I balance, suggesting a brief 20-min stimulation may be insufficient to induce an enhancement of global inhibitory activity (Fig. 2C). A 14-day multi-session tFUS (1 h/day) was then implemented, and the change in E/I balance was assessed by resting-state EEG recordings obtained the day after the final stimulation session (Fig. 2D). The multi-session tFUS resulted in a significant increase in the aperiodic exponent in female SCD mice, indicating a shift toward increased inhibitory activity, approaching levels observed in wild-type controls (Fig. 2E and G). Interestingly, an additional 20-min tFUS following the 14-day stimulation further significantly increased the aperiodic exponent compared to single-session tFUS in SCD mice, suggesting an enhanced network-level inhibitory effect potentially driven by synaptic plastic changes (Fig. 2F and I). For comparison, we further found that optogenetic activation of PV interneurons leads to a robust increase in the aperiodic exponent, mirroring the strengthened inhibitory function induced by multi-session tFUS, whereas activating SST interneurons significantly decreases the aperiodic exponent (Supplementary Fig. 2). These findings support the hypothesis that PV-mediated, rather than SST-driven, inhibition underlines the electrophysiological effects of tFUS.

### 3.3 Multi-session tFUS enhanced local expressions of GAD67 and PV interneurons

As repeated tFUS stimulations induced a significant and sustained increase in global inhibitory electrophysiological activity, we investigated cellular changes after multi-session 40 Hz PRF tFUS using IHC analysis and blinded assessment (Fig. 3A). Assessment revealed significantly higher GAD67-positive cells in the stimulated region compared to the contralateral side (Fig. 3B-C), whereas no such difference was observed in the sham-treated group (Supplementary Fig. 3A). These findings suggest that tFUS enhances inhibitory GABAergic signaling by increasing the capacity for GABA synthesis, as reflected by elevated levels of GAD67.

**Figure 3.**
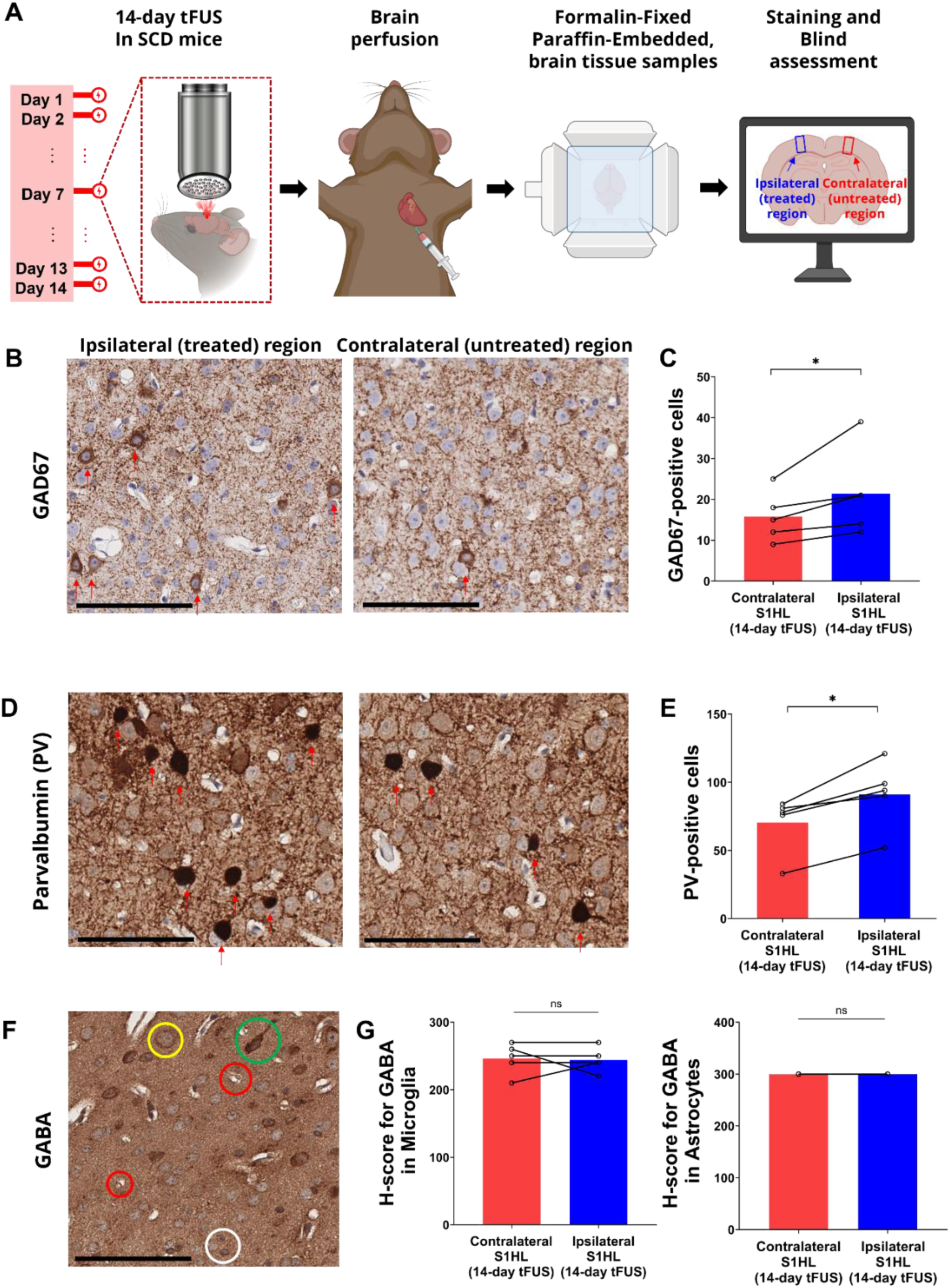
Multi-session tFUS increased neuronal GABAergic capacity assessed by IHC. (A) Schematic of histology processing workflow: mice underwent 14 days of tFUS, followed by brain perfusion, fixation, paraffin embedding, and blind IHC analysis comparing ipsilateral (stimulated) and contralateral (unstimulated) brain regions in SCD mice. (B) Representative IHC image of GAD67. (C) Quantification of GAD67-positive cells revealed a significant increase in the treated hemisphere following multi-session tFUS, suggesting enhanced capacity for GABA synthesis (n=5). (D) Representative IHC image of PV-positive neurons. (E) Quantification of PV-positive cells showed a significant increase in the stimulated hemisphere, indicating enhanced engagement of fast-spiking inhibitory interneurons (n=5). (F) Representative IHC image of GABA expression. Yellow, green, red, and white circles represent neuron, strong astrocyte, weaker oligodendrocyte, and microglial cells, respectively. (G) H-score quantification of GABA immunoreactivity in microglia and astrocytes showed no significant difference between ipsilateral and contralateral hemispheres, indicating that tFUS did not alter non-neuronal inhibitory signaling (n=5). \**p* <0.05 using *t* test with Wilcoxon matched-pairs signed rank test; ns, not significant. Scale bars represent 100 µm.

Building on the observed upregulation of GAD67, we next addressed the question of whether tFUS could modulate specific subtypes of GABAergic interneurons, particularly PV-expressing neurons, based on our findings related to theta oscillations, distinct E/I balance patterns, and prior evidence involving optogenetic inactivation of PV neurons [48]. Given their important role in maintaining E/I balance and known vulnerability in various neurological conditions [45], we sought to determine whether PV interneurons are upregulated following multi-session tFUS stimulation. Stimulation of left S1HL significantly increased the number of PV-positive cells at the site relative to the contralateral region (Fig. 3D-E), but not from sham treatment (Supplementary Fig. 3B). These findings suggest that tFUS strengthens local inhibitory networks by promoting activity or expression of the PV GABAergic neurons.

We further examined whether non-neuronal cell types such as microglia and astrocytes might also contribute. Since GABA is widely expressed in the brain, we alternatively used the H-score, a semi-quantitative method for evaluating protein expression [52,53], to compare GABA immunoreactivity between ipsilateral and contralateral regions. H-score analysis revealed that neither tFUS nor sham treatment led to significant changes in GABA expression within microglia and astrocytes across hemispheres (Fig. 3F-G and Supplementary Fig. 3C-D). Additionally, Iba1 levels remained unchanged following tFUS stimulation (Supplementary Fig. 3E-F), indicating a lack of glial activation. These results suggest that tFUS at 40 Hz modulates GABAergic signaling primarily through neuron-specific pathways in the humanized SCD mouse model.

### 3.4 Multi-dimensional histological analyses confirmed safety of multi-session tFUS

To facilitate potential clinical translation, we conducted a thorough safety evaluation by multi-dimensional histological analyses including H&E, TUNEL, and CD31 immunostaining, which are well-established approaches to evaluate tissue morphology [54], cellular integrity [55], and vascular architecture [56], respectively. Following multi-session tFUS stimulation, H&E staining revealed no gross morphological abnormalities, such as significant infiltration of macrophages or neutrophils, inflammation, necrosis, apoptosis, hemorrhage, fibrosis, vacuolation, or brain damage in either hemisphere under tFUS relative to the sham-treated group (Fig. 4A and Supplementary Fig. 4A), reinforcing the structural safety of the protocol. Additionally, TUNEL staining showed no evidence of apoptotic cell death as no TUNEL-positive nuclei were detected in either brain region of tFUS-treated or sham-treated animals (Fig. 4B and Supplementary Fig. 4B). Considering that vascular structures in the SCD mouse model may exhibit heightened vulnerability to mechanical perturbations^12^, we further examined vascular integrity using CD31, a widely accepted marker of endothelial cells. CD31 immunostaining demonstrated similar vascular distribution without signs of pathological angiogenesis, vessel disruption, or hemorrhage in both hemispheres from tFUS-treated and sham-treated groups (Fig. 4C and Supplementary Fig. 4C). Taken together, these results suggest that multi-session tFUS with a PRF of 40 Hz does not induce detectable cytotoxic effects or vascular damage in the brain samples used in the IHC analysis.

**Figure 4.**
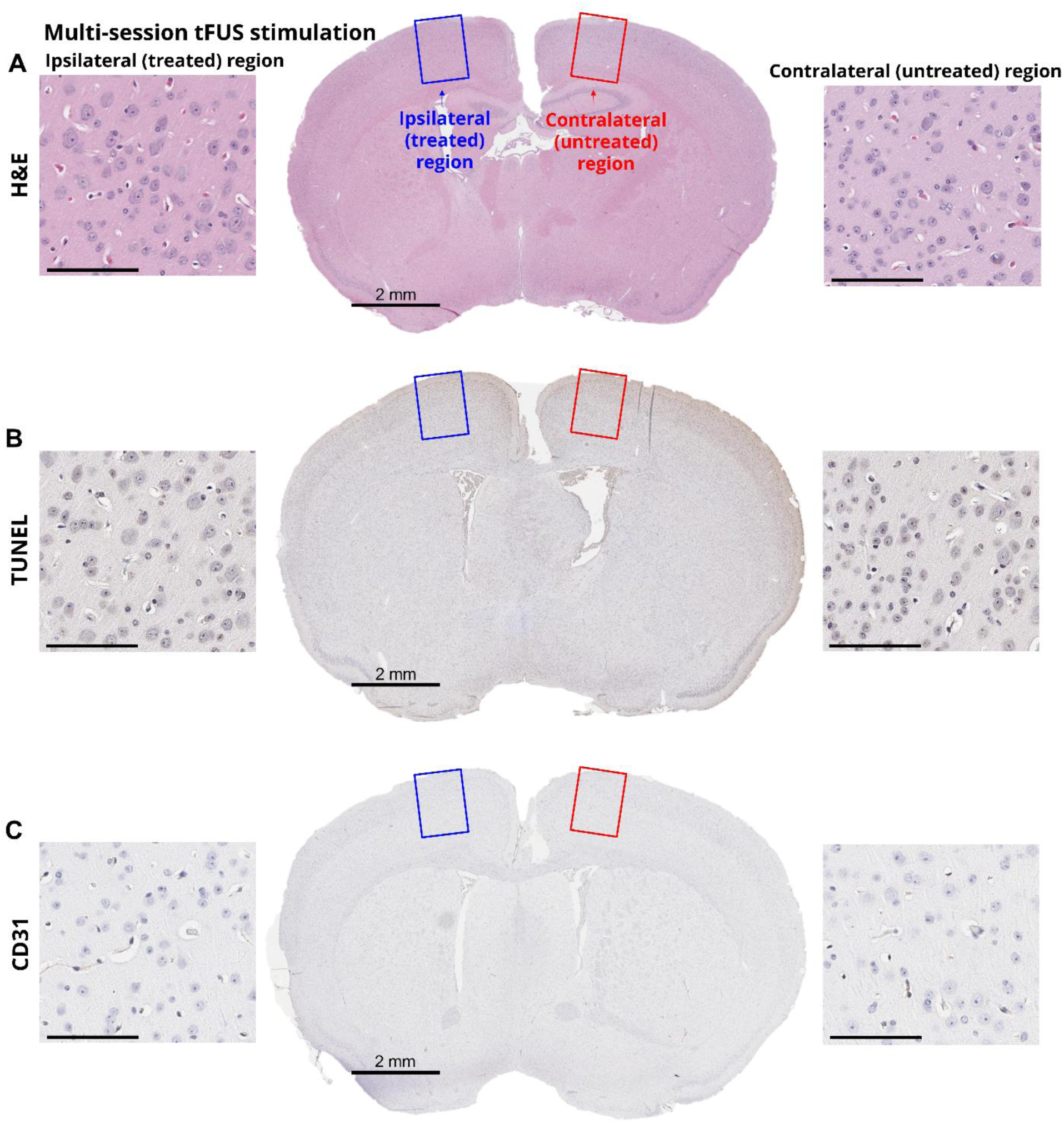
Multi-session tFUS did not induce pathological changes as assessed by histological analysis. (A) Representative H&E-stained section showed preserved tissue architecture in both the ipsilateral and contralateral brain regions following 14-day tFUS. No signs of macrophages, neutrophils, inflammation, necrosis, apoptosis, hemorrhage, fibrosis, vacuolation were detected across hemispheres. (B) Representative image of TUNEL staining revealed an absence of TUNEL-positive nuclei, indicating no evidence of necrosis or apoptotic cell death in either hemisphere. (C) Representative image of CD31 immunostaining indicated normal vascular architecture and distribution, with no signs of hemorrhage or abnormal angiogenesis. Scale bars represent 100 µm.

## DISCUSSION

Prior studies have highlighted the potential of tFUS in pain management. However, how tFUS mediates its effects is still poorly understood. In this study, we have demonstrated that low-intensity tFUS with a PRF of 40 Hz engaged PV GABAergic interneurons in a humanized mouse model of chronic pain using combined electrophysiological and molecular approaches. These included enhanced local and global inhibitory activity in the pain-processing brain circuit, as indicated by suppression of low-frequency oscillations at S1HL, elevated aperiodic exponent derived from resting-state EEG, and increased GAD67-immunoreactivity (i.r.) and PV interneuron-i.r. in the stimulated region. Compared to high-intensity tFUS used in ablative protocols, which eliminate cellular activity in the targeted area but risk collateral damage to adjacent healthy circuits potentially leading to adverse side effects [57], this low-intensity tFUS enables targeted neuromodulation while preserving structural and functional integrity. Importantly, we showed tFUS effects without evidence of glial activation, cellular apoptosis, or vascular disruption, confirming safety and neuron-specificity. This specificity underscores its potential as a non-invasive and precise neuromodulation approach for chronic pain management, paving the way for promising therapeutic applications.

Our results demonstrated that 40 Hz tFUS led to upregulation of both GAD67 and PV expressions, two molecular hallmarks of the GABAergic inhibitory system. Given prior evidence that GAD67 deficiency in PV interneurons resulted in impaired inhibitory transmission and cortical disinhibition [39,58], the observed post-tFUS upregulation supports the restoration of inhibitory activity. GAD67 is the enzyme that catalyzes synthesis of the inhibitory neurotransmitter GABA from glutamate [36], and is currently being explored as a therapeutic target for treating pain, including via gene therapy approaches [59,60]. Recent study in SCD patients shown elevated brain glutamate, potentially suggesting the correlation between the excitatory neurotransmitter and pain hypersensitivity [61]; thus, increased GAD67 expression may help counterbalance this excitatory drive by enhancing the glutamate to GABA conversion. Our approach utilizing non-invasive tFUS led to upregulation of GAD67, supporting its potential as a convenient and cell-targeted strategy for pain relief. In parallel, GABAergic neurons in S1 primarily consist of PV, SST, and vasoactive intestinal peptide (VIP) interneurons [62]. These inhibitory neuron subtypes differ in distribution, firing properties, and synaptic targets [63]. Among them, PV interneurons are the dominant GABAergic population, characterized by rapid, high-frequency spiking and strong perisomatic inhibition. Their loss significantly heightens pain sensitivity [64–66], highlighting the relevance of PV GABAergic neurons for effective pain treatment. In our study, we observed a significant increase in PV-positive cell expression in the tFUS-stimulated area, positioning tFUS as a cell-type specific modality in this context.

A holistic interpretation of both electrophysiological and IHC findings revealed two complementary clinical applications for the use of tFUS with a PRF of 40 Hz. A single-session tFUS induced suppression of low-frequency EEG rhythms, consistent with short-term circuit inhibition, but did not alter global E/I balance. Thus, it appeared effective for providing rapid relief from acute or tonic pain, which typically arose from inflammation, healing, or mild injury and is generally reversible. It modulated cortical excitability locally and temporarily without inducing detectable morphological changes. As a non-pharmacological alternative, tFUS holds particular promise for patients who may be contraindicated for pharmacological analgesics, which are associated with risks such as gastrointestinal distress, organ toxicity, dependency, and overdose [1–3,67,68]. In contrast, multi-session tFUS accumulated over time, engaging inhibitory circuits through potential synaptic plasticity rather than short-term modulation. These findings aligned with prior results from repetitive TMS, where multiple TMS stimulation induced inhibitory plasticity in cortical networks [69]. This adaptive reorganization of a previously dysregulated inhibitory circuit highlighted a novel therapeutic mechanism of low-PRF tFUS, complementing previously studied pulse-locked neuromodulation or mechanosensitive ion channel effects [70,71]. By showing elevated PV-i.r. and GAD67-i.r. alongside persistent electrophysiological changes, our study provided post-stimulation mechanistic evidence of functional network remodeling. While previous reports have focused on acute electrophysiological responses, behavioral outcomes, or self-reported pain ratings [28–33], our findings expanded the mechanistic landscape to include molecular and cellular adaptations in inhibitory circuits driven by repeated subthreshold entrainment of PV neurons. Additionally, when single-session tFUS was applied to 14-day-treated SCD mice, the stimulation effect was significantly greater compared to that observed with single-session tFUS in naïve SCD mice, indicating that non-invasively repetitive stimulations can drive synaptic plasticity even when each session is subtle and transient. Importantly, we observed no signs of cellular or vascular damage across the 14-day stimulation compared to controls, which is an essential safety consideration for any long-term neuromodulation therapy. These results positioned low-intensity tFUS with 40 Hz PRF as a versatile and precision intervention, with flexible dosing paradigms tailored for different clinical needs: single-session use for episodic pain control, and multi-session delivery for chronic pain alleviation through inhibitory circuit restoration. Moreover, the combined use of electrophysiological and immunohistochemical endpoints provided cross-validated evidence for a reliable, non-invasive biomarker with translational relevance. The power spectral changes and the aperiodic exponent derived from EEG, a clinically accessible modality, closely reflected shifts in cortical inhibitory activity and mirrored molecular findings of GAD67 and PV-positive interneurons upregulation. This convergence of readouts suggested that EEG-based measurements may serve as practical surrogate biomarkers for monitoring neuromodulatory engagement in clinical settings.

While our study provided multi-level evidence that repeated 40 Hz tFUS strengthened cortical inhibitory activity and promoted long term circuit reorganization, several directions remained to be explored. First, although the use of PV and GAD67 markers was deliberate and mechanistically informative, future studies could incorporate a broader panel of IHC markers to map the GABAergic and Glutamatergic responses more comprehensively. For example, the inclusion of SST, VIP, CaMKII, and GABA receptors could allow for a more detailed dissection of cell type and synapse specific changes. Our results aligned with the overall low-frequency power suppression observed during optogenetic activation of PV neurons, the statistical significance inconsistency in the beta band could suggest other potential mechanisms contributing to the inhibition, such as the activation of other GABAergic interneurons or the inhibition of excitatory glutamatergic neurons. Mapping the involvement of additional GABAergic subtypes, including SST, VIP, and their synaptic interactions with excitatory neurons would greatly enhance our understanding of circuit-level reorganization and the precision of tFUS neuromodulation, given the distinct roles these subtypes contribute to the overall sensory perception [72]. Second, when grand-averaging EEG responses across 17 mice, we observed a relatively broader distribution of activity (Supplementary Fig. 5), likely reflecting inter-animal anatomical variability including slight differences in electrode placement, skull thickness, and cortical geometry, which can influence the precise localization of evoked responses. Although this variability is less critical given the relatively thin mouse skull, future studies should consider individualized targeting strategies or improved anatomical registration approaches to better align responses across subjects. Lastly, the translational pathway from rodent models to human application required careful calibration of stimulation parameters including acoustic pressures and stimulation pulse paradigm. In this study, we used humanized SCD mouse model; however, careful consideration must be given to the selection of stimulation parameters due to fundamental biological differences between rodents and humans, including differences in cortical organization and interneuron diversity [73]. Computational modeling, paired with validation in large animal models, will be critical to defining safe and effective stimulation protocols for human use.

Altogether, our findings highlighted that tFUS engages PV GABAergic inhibitory circuits to counteract chronic pain features, supported by electrophysiological and cellular analyses. These insights positioned tFUS neuromodulation as a clinically meaningful avenue for restoring inhibitory balance in the brain, with the potential to redefine therapeutic strategies for chronic pain and disorders rooted in circuit dysfunction.

## Data availability

The data supporting the conclusions of this study are provided in the paper and supplementary materials. Additional data will be made available at a public repository when the paper is accepted.

## CRediT authorship contribution statement

**Min Gon Kim:** Conceptualization, Methodology, Investigation, Formal Analysis, Writing – Original Draft, Writing – Review & Editing. **Chih-Yu Yeh**: Conceptualization, Methodology, Investigation, Analysis, Writing – Original Draft, Writing – Review & Editing. **Huan Gao**: Methodology, Investigation, Writing – Review & Editing. **Keunhyung Lee**: Methodology, Writing – Review & Editing. **Kalpna Gupta:** Methodology, Resources, Writing – Review & Editing. **Bin He**: Conceptualization, Methodology, Investigation, Supervision, Resources, Writing – Review & Editing.

## Acknowledgments

This work was supported in part by NIH (Grant Nos. NS131069, NS124564, EB029354, HL147562, CA263806, and AT012868). The authors thank Dr. Kai Yu for useful discussions, Dr. Julie Feldstein from Hitowiz company for providing the histologic analysis, and Dr. Donovan Argueta for preparing paraffin-embedded brain specimens. Some cartoons in figures and supplementary figures were created with BioRender.com.

